# Pan-cancer evolution signatures link clonal expansion to dynamic changes in the tumour immune microenvironment

**DOI:** 10.1101/2023.10.12.560630

**Authors:** Xinyu Yang, Wei Liu, Geoff Macintyre, Peter Van Loo, Florian Markowetz, Peter Bailey, Ke Yuan

## Abstract

Cancer is an evolutionary process characterised by profound intra-tumour heterogeneity. Intra-tumour heterogeneity can be quantified using in silico estimates of cancer cell fractions of tumour-specific somatic mutations. Here we demonstrate a data-driven approach that uses cancer cell fraction distributions to identify 4 robust pan-cancer evolutionary signatures from an analysis of 4,146 individual tumour samples (TCGA) representing 17 distinct cancer types. Evolutionary signatures defined a continuum of cancer cell fractions representing neutral evolution, clonal expansion and fixation. Correlation of evolutionary signatures with programs representing distinct mutational and biological processes demonstrated that individual tumours enriched for clonal expansions and fixations were associated with immune evasion and distinct changes in the tumour immune microenvironment. We observed a dynamic switch between adaptive and innate immune processes as tumours undergo clonal fixation and escape immune surveillance. We also identify mutational processes underpinning different modes of tumour evolution and demonstrate that switching between adaptive and innate immune cell populations is accompanied by the clonal expansion of driver genes that modulate tumour-stroma interactions^1^.

## Introduction

Genetic intra-tumour heterogeneity (ITH) has recently emerged as a universal feature of tumours^2, 3^. ITH arises through an evolutionary process involving both cancer cells and the tumour microenvironment^4^ providing selective cellular adaptation that supports the fitness of evolving tumour clones^5, 6^. Large-scale bulk sequencing efforts of cancer genomes^7–10^ have revealed tens of millions of genomic alterations, providing an invaluable resource for understanding the complexities of cancer evolution. While studies have focused on inferring evolutionary dynamics from ITH^11–16^, it is largely unknown how ITH relates to the tumour microenvironment.

Tumours evolve by the selection of specific traits that provide a survival advantage^17–19^. Adaptation to diverse pressures including the host’s immune system and cytotoxic stress shape the evolution of cancer cells by driving selective genomic modifications^20^. Accumulation of mutations that overcome these selection pressures allows cancer cells to populate most of the tumour and escape the immune system^21^. Dynamic shifts in tumour cell-intrinsic and extrinsic selection pressures control cancer evolution^22^. Distinct tumour cell-extrinsic selection pressures shape different tumour types with immune adaptation and external carcinogens providing alternative evolutionary trajectories to field cancerization^23, 24^.

Each tumour is an independent evolution running its own course. A long promise of studying cancer evolution has always been finding commonalities in how tumours evolve such that the underlying driving forces of a malignancy can be characterised to a point where the dynamics of tumour progression can be accurately modelled. Analytical approaches to identify such commonalities have so far included phylogenetic trees^25^ and neutral and selection dynamics from predefined mechanistic models^12^. A challenge shared with these approaches is that it is not straightforward to associate ITH with biological hallmarks acquired during the multistep development of human tumours using the same patient cohort.

In this study, we propose a new machine learning framework to identify common patterns of cancer evolution dynamics, which we refer to as consensus evolutionary dynamic signatures (ES), which can bridge the gap between evolutionary analysis and cancer hallmarks, making an assessment of the consequences of ITH possible in bulk sequenced tumour samples.

## Results

### Modelling evolutionary dynamics in cancer genomes

Our method addresses two key limitations of the neutral formulation in Williams et al^12^. First, instead of modelling variant allele frequency (VAF), we model cancer cell fractions (CCFs). CCF represents the percentage of cancer cells bearing a mutation in a tumour sample. This change allows the framework to correct for normal contamination and copy number alterations, thereby, maximising the number of SNVs eligible for modelling and improving SNV distributions (histograms). Second, we introduce a generalised formulation to the neutral model to generate full CCF distributions. We achieve the transition from modelling the number of mutations (M) as a function of VAF to the number of mutations as a function of CCF using a change of variable.

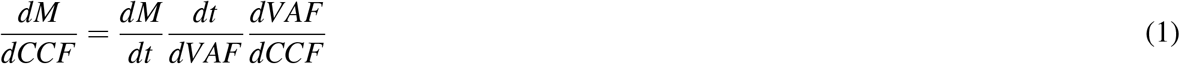

The generalised formulation is designed to capture various growth and population parameters for mutations with different growth behaviours in real tumours. Some mutations may follow a neutral growth pattern with a constant growth rate, while others exhibit selective growth with varying growth rates possibly.

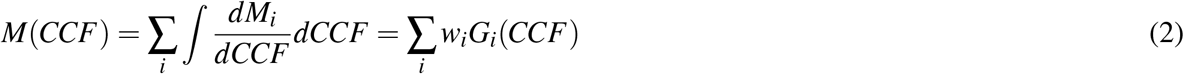

where *i* represents *i* mutation groups with different growth behaviours. The *G*_*i*_(*CCF*) functions represent growth functions describing these growth behaviours, and the parameters *w*_*i*_ represent how these functions give rise to the observed CCF distribution. Mutations with the same growth behaviours share the same growth functions and parameters, resulting in clusters that follow certain distributions. Without loss of generality, in the case of *G*(*CCF*) being an exponential growth function and *i* = 1, we can recover the neutral model^12^. More generally, cluster-like patterns in CCF distributions can be represented as a weighted sum of different growth functions, accommodating a broader range of growth behaviours beyond the specific case of exponential growth.

Analytically solving the equation is infeasible, as we do not know all the underlying growth functions that contribute to the model. However, by pooling samples in a cohort, we can have a data-driven solution to the above equation using matrix factorisation:

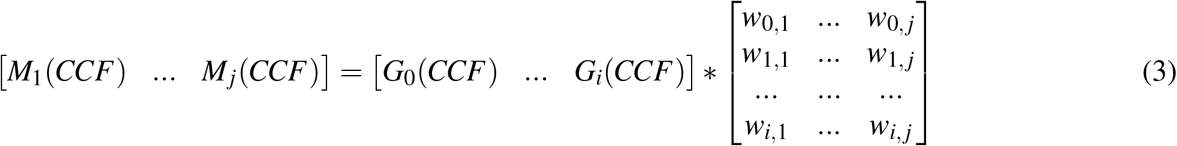

where *j* represents the *j*th sample and *G*_*i*_(*CCF*) represents the *i*th growth function shared across tumours, namely a signature. We use non-negative matrix factorization (NMF) to solve this matrix factorisation problem and simultaneously extract the signatures and contributions of each signature among samples. Similar methods have been successfully applied to mutational signatures^26, 27^ and copy number signatures^28–30^.

### Four consensus signatures of evolutionary dynamics in 2917 cancers

To identify robust ES, 2917 whole-exome sequencing samples across 12 cancer types from The Cancer Genome Atlas (TCGA)^31^ were qualified for inclusion in the analysis with the following criteria (Figure 1a): 1) patients with a minimum of 30 reliably called private mutations. 2) patients with a suggested average depth>120x^32^. 3) patients with high-quality cancer cell fraction estimation. These CCF estimations were obtained using CCube^33, 34^, which has been shown to be robust across several benchmarks^35, 36^. 4) cancer types with at least 100 samples. We constructed sample-by-CCF matrices for each cancer type. Each row of a sample-by-CCF matrix consists of the number of mutations that fall in 100 discretised bins between 0 and 1 over CCF, depicting the distribution of M(CCF) for each sample. We first performed NMF for sample-by-CCF matrices of each cancer type and thus obtained type-wise evolutionary dynamics signatures (Supplementary Figure 2). The optimal number of signatures for each cancer type was chosen by performing 1000 runs of the algorithm with different random seeds and 1,000-time shuffles of the input matrix to avoid over-fitting (Supplementary Figure 1). Unsupervised hierarchical clustering was then performed on all type-wise signatures, and the number of clusters was suggested by the Hubert index. The final set of evolutionary dynamics signatures was obtained by normalising and averaging type-wise signatures for each cluster (Figure 1b). Actually, we also observed similar signatures by directly performing NMF on pan-cancer datasets, further reinforcing the robustness of our method.

**Figure 1.**
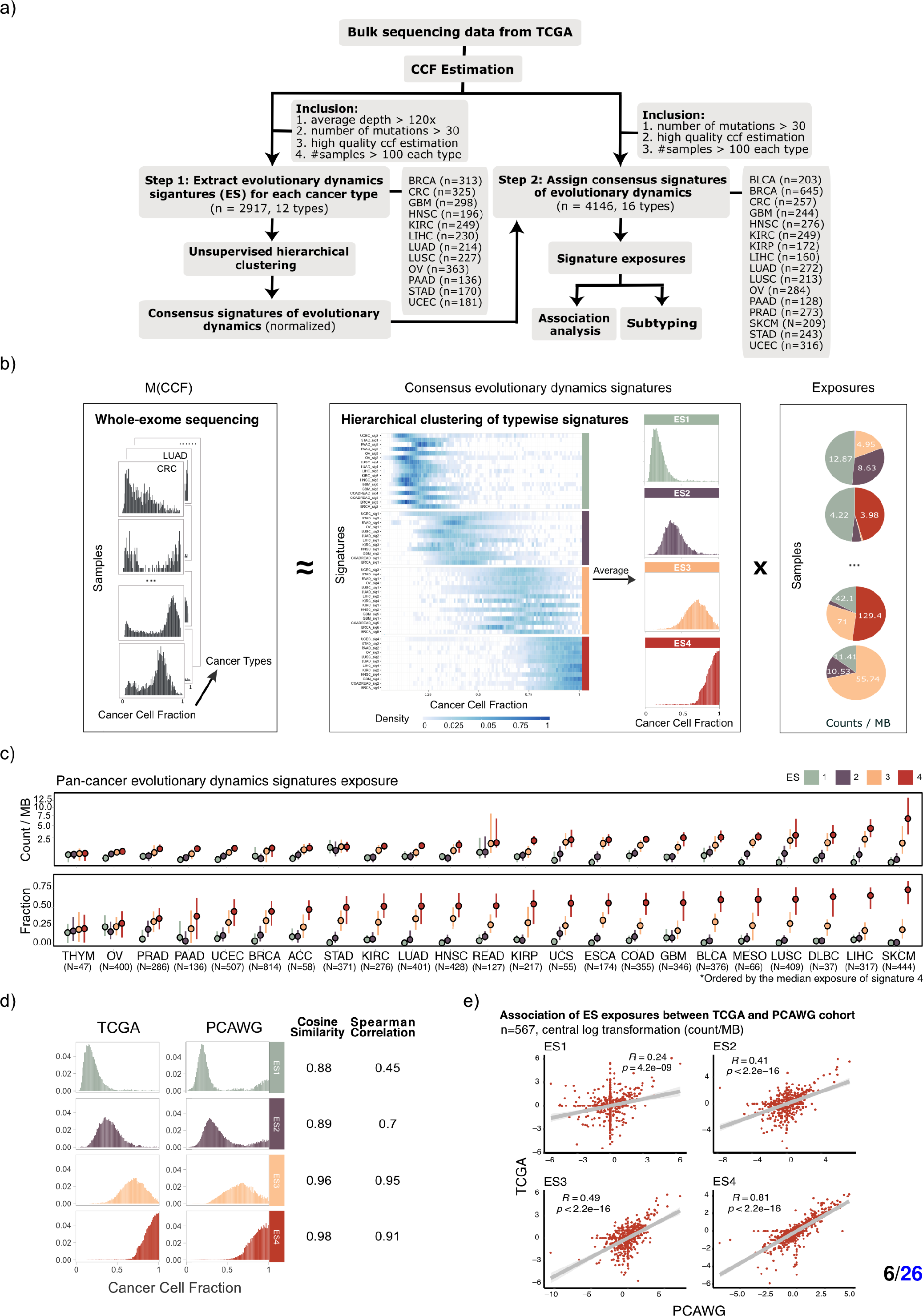
(Previous page.) Overview of study design and framework of identifying consensus evolutionary dynamics signatures. **a**, Flowchart of patient inclusion and downstream analysis. Two-step process of estimating patterns of evolutionary dynamics: 1) extracting consensus signature of evolutionary dynamics for each cancer type. 2) hierarchical clustering captures the most common patterns across cancer types. Downstream analyses were performed by estimating the exposure of each consensus signature of evolutionary dynamics for TCGA samples. **b**, Illustration of how non-negative matrix factorization (NMF) identifies the consensus signatures of evolutionary dynamics (ES). **c**, Estimation of contributions of ESs in a single tumour across cancer types. **d**, Identified ESs in two independent cohorts, TCGA (WES) and PCWAG (WGS). The cosine similarity and correlation coefficient between these cohorts are indicated for each signature and are provided in the Supplementary Table. **e**, Scatter plots depicting the correlation of ES exposure for the same patients between TCGA and PCAWG cohorts (n=567). Spearman correlations were estimated after applying a central log transformation to each signature exposure.

Our analysis identified four consensus evolutionary dynamics signatures using the TCGA cohorts (Figure 1b). Evolutionary Dynamics Signature 1 (ES1) displays a left-skewed distribution concentrated in the low CCF region, close to 0. ES1 appears to adhere to the 1/f distribution, which has previously been discussed in terms of its relation to neutral evolution^12, 37, 38^. ES2 exhibits a bell-shaped distribution, primarily concentrated within the CCF range of 0.25 to 0.55. ES2 could potentially represent mutations coming out of the long tail of a typical neutral peak. ES3 appears to have a bell-shaped distribution similar to ES2, yet it accommodates mutations shifting to higher CCF ranges, typically falling within the range of 0.6 to 0.8. This level of CCF indicates mutations getting close to being fixed in the population, demonstrating evidence of one or several subclones becoming the dominant clone in the sample. Tumours with strong exposure to ES3 could therefore be under active subclonal expansion. In comparison to ES3, ES4 displays a more pronounced shift towards a close to 100% CCF. It also exhibits a long-tail shape that accommodates mutations that gradually approach fixation (CCF=1). Strong ES4 exposure indicates the tumour has undergone significant clonal expansion.

Estimation of contributions of ES, namely signature exposure, can provide a coarse estimation of evolutionary dynamics in the unit of tumour mutation burden (Count/MB) in a single tumour. This estimation allows for downstream analysis to further refine the definitions and interpretation of ESs in terms of DNA damage status, immune landscape, biological process, and clinical relevance. Here, we assigned these four evolutionary dynamics signatures on 4146 samples across 16 cancer types to estimate the contribution of each signature in each sample (Figure 1c). These 4146 samples were included based on criteria similar to those used for signature identification but with relatively more leniency, as they did not require an average depth >120x (Figure 1a).

Whole-genome sequencing dataset from the Pan-Cancer Analysis for Whole Genomes (PCAWG)^10^ was included for validation. We found highly similar evolutionary dynamics signatures within the TCGA and PCAWG cohorts, especially for ES3 and ES4 (ES3: cosine similarity = 0.96, Spearman correlation = 0.95; ES4: cosine similarity = 0.98, Spearman correlation = 0.91; Figure 1d). These observations demonstrate the robustness of both our analytical methodology and ESs across whole-exome sequencing and whole-genome sequencing. We further associated ES exposures among 567 patients concurrently sourced from both the TCGA and PCAWG cohorts. We observed a strong association within ES4, followed by ES2 and ES3 (ES2: r=0.41, *P* < 2.2*e* −16; ES3: r=0.49, *P* < 2.2*e* −16; ES4: r=0.81, *P* < 2.2*e* −16; Figure 1e) between ES exposures. ES1 displayed the lowest level of correlation, possibly attributed to limited coverage to detect low-frequency mutations within the tumour population using whole-genome sequencing.

### Signatures associated with DNA damage and biological processes

During cancer evolution, genomic instability provides materials for selection and favours tumour progression through multiple biological processes^39, 40^. To systematically investigate the underlying biological process and DNA damage related to evolutionary dynamics signatures, we correlated ES exposures with factors related to DNA damage from^41^, including copy number burden^42^, homologous recombination deficiency (HRD)^42^, intra-tumour heterogeneity (ITH)^43^, aneuploidy score^43^, predicted neoantigen^41^ and mutation rate^41^. We also retrieved weights of COSMIC mutational signatures for TCGA patients, which are characterised by mutations arising from specific mutagenesis processes such as DNA replication infidelity, exogenous and endogenous genotoxin exposures, defective DNA repair pathways, and DNA enzymatic editing^27^.

Our results suggest that ES1 is generally negatively associated with DNA damage (including copy number variation burden, HRD and Aneuploidy Scores), predicted neoantigens, mutation rate (Figure 2c) and only a few mutational processes like mitotic clock process and dMMR process (Figure 2d). For example, ES1 negatively correlated with HRD in BLCA (*r* = −0.47, *P* = 7 × 10^−5^) and LUSC (*r* = −0.45, *P* = 2 × 10^−5^), copy number burden score "Fraction Altered"in LUAD (*r* = −0.51, *P* = 2 × 10^−7^) and PAAD (*r* = −0.45, *P* = 6 × 10^−4^), and SNV neoantigen in LUAD (*r* = −0.31, *P* = 3 × 10^−3^) and LUSC (*r* = −0.37, *P* = 7 × 10^−4^) (Figure 2a). ES1 exhibited a positive association with the mitotic clock process in CRC, STAD, and UCEC (*r* = −0.64, *P* = 3 × 10^−31^). ES1 was also associated with important cancer-driven mutational processes, such as the APOBEC process in HNSC and BRCA, as well as the dMMR process in UCEC (*SBS*15 : *r* = 0.58, *P <* 1 × 10^−4^) (Figure 2b).

**Figure 2.**
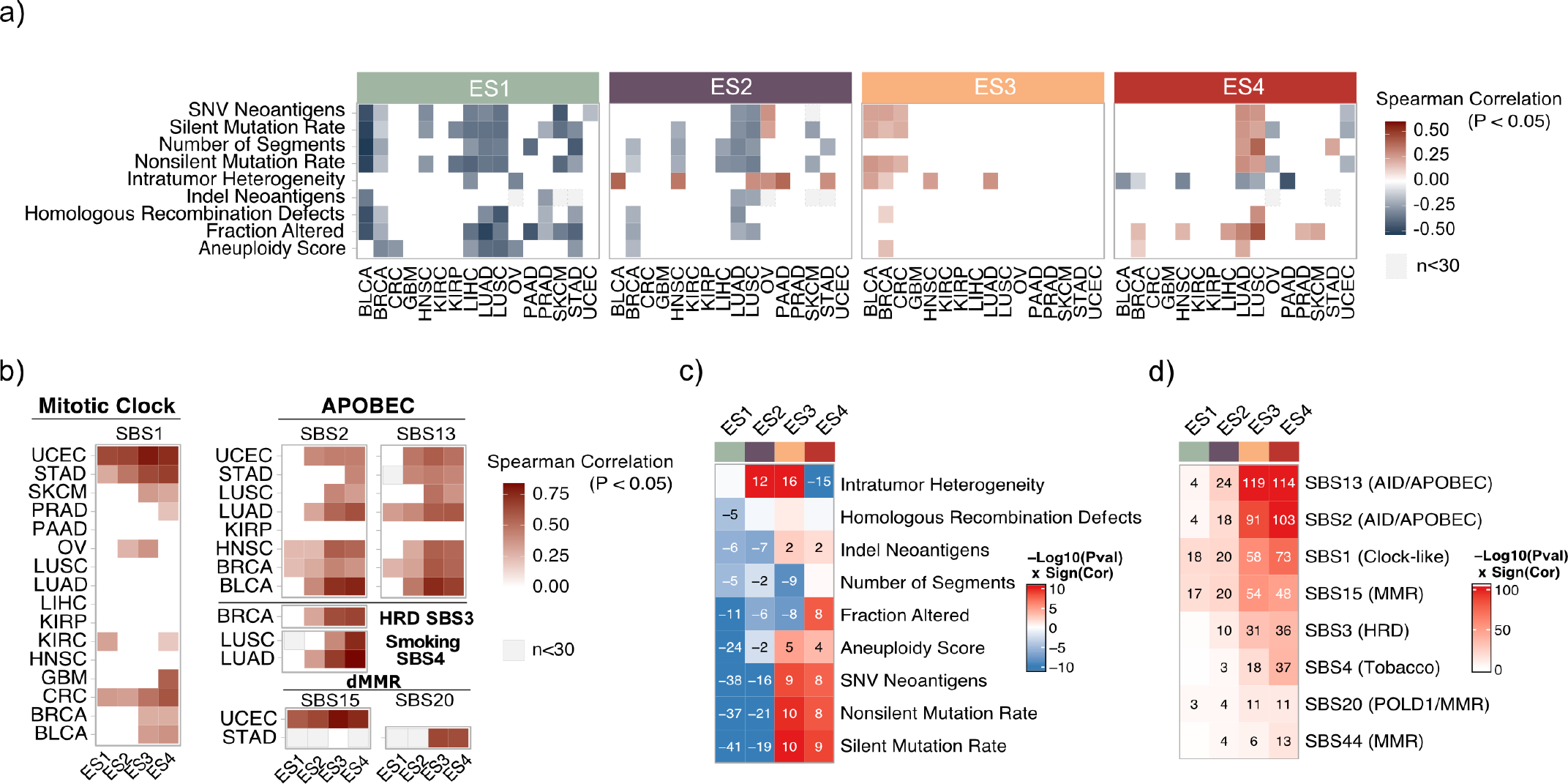
Underlying biological process and molecular characterisation behind the evolutionary dynamics signatures. **a**, Pan-cancer association between ES exposures and copy number burden, HRD, ITH, DNA damages scores, predicted neoantigen and mutation rate in TCGA (only associations with a false-discovery rate *P* < 0.05 and at least 30 samples are shown). **b**, Pan-cancer association between ES exposures and mutational signatures. Spearman correlation coefficients and adjusted *P* values are as indicated (only associations with a false-discovery rate *P < 0*.*05* are shown). **c-d**, Association between ES exposures and copy number burden, HRD, ITH, DNA damages scores, predicted neoantigen, mutation rate and mutational signatures in all TCGA samples. Spearman correlation and adjusted P values are as indicated (only associations with a false-discovery rate *P* < 0.05 are shown).

In comparison to ES1, ES2 showed similar but slightly weaker associations with DNA damage scores across cancer types. Additionally, ES2 was positively correlated with the ITH score in several cancer types, such as HNSC (*r* = 0.34, *P* = 3 × 10^−4^) and PAAD (*r* = 0.38, *P <* 0.05) (Figure 2a). Furthermore, similar to ES1, ES2 displayed a positive association with the Mitotic Clock, APOBEC, and dMMR processes. However, this association was present in more tumour types for ES2 than ES1 (Figure 2b, d).

In general, ES3 and ES4 were both positively associated, opposite to ES1 and ES2, with aneuploidy score, neoantigens, and traditional features of selection, such as silent/non-silent mutation rates (Figure 2c). They also made a significant contribution to APOBEC, MMR, and HRD processes (Figure 2d). For example, positive associations were observed between ES4 and APOBEC SBS2 (BLCA: *r* = 0.77, *P* = 1 × 10^−35^; LUAD: *r* = 0.62, *P* = 9 × 10^−16^), HRD SBS3 (BRCA: *r* = 0.66, *P* = 2 × 10^−26^), smoking SBS4 (LUAD: *r* = 0.84, *P* = 2 × 10^−27^; LUSC: *r* = 0.75, *P* = 2 × 10^−10^) and dMMR SBS15 (UCEC: *r* = 0.75, *P* = 3 × 10^−7^) (Figure 2b). Interestingly, we observed an overall opposite pattern in the association of ES3 and ES4 with ITH and copy number burden score” Fraction Altered”, suggesting that distinct states in the selection are captured, respectively (Figure 2c). For example, ES4 positively correlated with “Fraction Altered” in LUSC (*r* = 0.44, *P* = 6 × 10^−9^), whereas ES3 was negatively correlated (*r* = −0.5, *P* = 7 × 10^−11^). ES4 was negatively associated with ITH, opposite to ES3, in HNSC (ES3: *r* = 0.24, *P* = 8 × 10^−3^; ES4: *r* = −0.32, *P* = 4 × 10^−5^) and LUAD (ES3: *r* = 0.27, *P* = 3 × 10^−4^; ES4: *r* = −0.28, *P* = 2 × 10^−4^) (Figure 2a).

### Signatures associated with immune infiltration

The immune microenvironment influences tumour evolution in terms of the complex interplay between cancer cells and infiltrating immune cells and can be both prognostic and predictive of response to immunotherapy^22, 41, 44^. The immune tumour microenvironment (TME) across cancer types can be characterized by various immunogenomics methods, including assessment of total lymphocytic infiltrate from genomic and H&E image data, immune cell fraction from deconvolution analysis of mRNA-seq data and immune gene expression signatures^41^. Here, we systematically investigated the relationship of ESs with these factors relating to the TME.

Proliferation and wound healing signatures were related to the cell cycle phase and associated with poor prognosis in cancer patients^45, 46^. We found that ES1 and ES2 were generally negatively associated with proliferation and wound healing (Figure 3b). Specifically, ES1 was negatively associated with wound healing and proliferation in BLCA, LIHC, LUAD, and LUSC (Wound Healing: *r* = −0.26, *P* = 2 × 10^−2^; Proliferation: *r* = −0.33, *P* = 3 × 10^−3^), STAD and UCEC. Interestingly, ES4 was positively associated with wound healing (*r* = 0.34, *P* = 3 × 10^−5^) and proliferation (*r* = 0.3, *P* = 3 × 10^−4^) in LUSC, whereas ES1, ES2 and ES3 (Wound Healing: *r* = −0.32, *P* = 1 × 10^−4^; Proliferation: *r* = −0.31, *P* = 2 × 10^−4^) were all negatively associated (Figure 3a).

**Figure 3.**
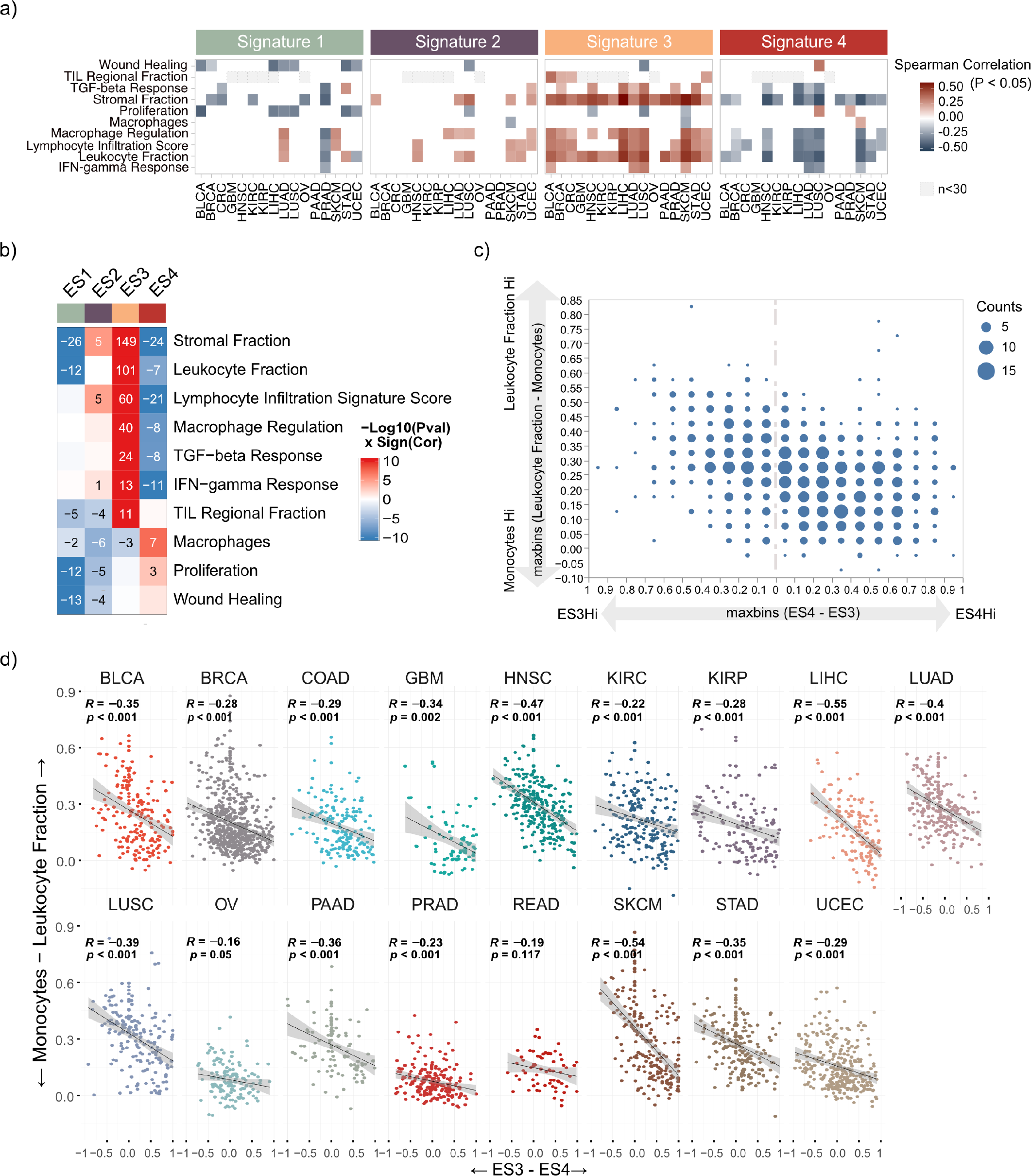
tumour microenvironment associated with deviance between ES3 and ES4. **a**, Pan-cancer association between ES exposures and immune signatures in TCGA (Only associations with a false-discovery rate *P* < 0.05 and at least 30 samples are shown). **b**, Association between ES exposures and immune signatures in TCGA. Spearman correlation coefficients and adjusted P values are as indicated (only associations with a false-discovery rate *P* < 0.05 are shown). **c**, Scatter plot showing a general negative association between immune infiltration (leukocyte fraction minus monocytes fraction) and deviance between ES3 and ES4 in TCGA. Blue dots denote the intensity of points overlapped. **d**, Scatter plots showing negative associations between immune infiltration (leukocyte fraction-monocytes fraction) and deviance between ES3 and ES4 across cancer types.

In general, we observed ES3 was positively associated with immune infiltrations (including stromal fractions, leukocyte fractions, lymphocyte infiltration scores, TGF-*β* response, IFN-*γ* response and macrophage regulation), opposite to ES4 (Figure 3b). Specifically, ES3 was strongly positively associated with the leukocyte fraction in most types of cancer (HNSC: *r* = 0.4, *P* = 4 × 10^−9^; LIHC: *r* = 0.52, *P* = 6 × 10^−8^; LUAD: *r* = 0.39, *P* = 2 × 10^−7^; LUSC: *r* = 0.48, *P* = 3 × 10^−10^; SKCM: *r* = 0.51, *P* = 2 × 10^−10^), whereas ES4 was negatively associated with the leukocyte fraction (HNSC: *r* = −0.42, *P* = 5 × 10^−10^; LIHC: *r* = −0.38, *P* = 9 × 10^−5^; *LUAD* : *r* = −0.37, *P* = 6 × 10^−7^; LUSC: *r* = −0.44, *P* = 1 × 10^−8^; SKCM: *r* = −0.41, *P* = 3 × 10^−7^) (Figure 3a).

The extensive observed opposing associations of ES3 and ES4 with leukocyte fraction prompted us to investigate further the relationship between adaptive immune response represented by leukocyte fraction and innate immune response represented by monocytes, and differences between ES3 and ES4. We found that high ES4 was associated with increased monocyte signature enrichment, while high ES3 was related to increased leukocyte fractions across cancer types (Figure 3c, d), which suggests a transition from adaptive immune response to innate immune response with the increase of ES4 proportion.

### Evolutionary dynamics subtypes reflect distinct immune mechanisms during cancer evolution

Given the results from previous analyses, which have demonstrated distinct states and immune mechanisms associated with ES3 and ES4 in the context of cancer evolution, we characterised differences in cancer hallmarks between ES3 and ES4. Specifically, we focused our investigation on colorectal cancer (CRC), stomach adenocarcinoma (STAD), and uterine corpus endometrial carcinoma (UCEC), which have been previously studied regarding mutator phenotypes (POLE, MSS, MRR) and underlying immune escape mechanisms^22^.

Our findings reveal that the mean ES3 proportion is significantly higher in tumours exhibiting evidence of immune escape compared to those without such evidence (*P* = 1 × 10^−4^, Kruskal-Wallis’s test). In contrast, the distribution of ES4 proportion does not show a significant difference between the immune escape groups (**Figure 4d**). Moreover, we observe that the mean difference between ES3 and ES4 proportions is lower in MSS tumours compared to those with MMR and POLE mutations (*P* = 4 × 10^−7^, Kruskal-Wallis test). Additionally, the mean difference between monocytes and leukocyte fraction in MSS tumours is lower compared to tumours with MMR and POLE mutations (*P* = 8 × 10^−7^, Kruskal-Wallis test), as depicted in Figure 4e.

**Figure 4.**
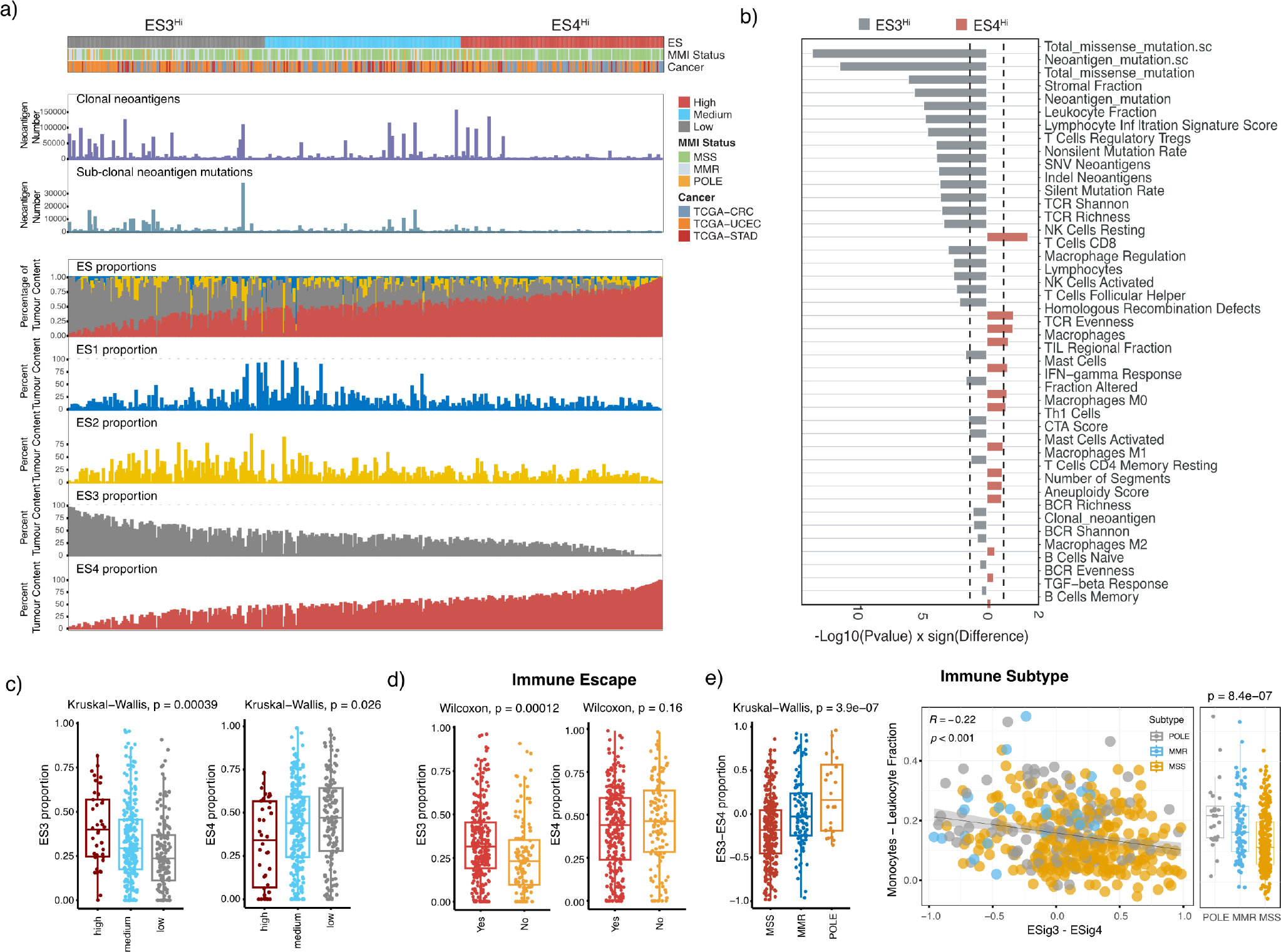
Survey of evolutionary dynamics subtypes in CRC, UCEC, STAD. **a**, Overview of ESs distribution over evolutionary dynamics subtypes. Samples were categorized into three groups (high, medium, low) through a trisection based on the difference between ES3 and ES4 proportions, where group high and low are defined as ES3^Hi^ and ES4^Hi^ in the following analyses. **b**, Differences in immune infiltration were compared between ES subtypes and p values were shown by Wilcoxon signed-rank test. **c**, Boxplots show the distribution of ES3 and ES4 proportions stratified between three groups (high, medium, low) and are annotated by a Kruskall-Wallis P value. **d**, Boxplots show the difference of immune escape scores stratified between ES subtypes and are annotated by a Kruskall-Wallis P value. **e**, Immune infiltration distribution over ES3 and ES4 (ES3-ES4) deviation with a Spearman correlation coefficient and P value. Boxplots show the difference in immune infiltration across MSI status and are annotated by a Kruskall-Wallis P value.

To elucidate the states of evolutionary dynamics characterized by distinct immune mechanisms, we categorised tumours into ES3^Hi^ or ES4^Hi^ subtypes, based on the differences in ES3 and ES4 proportions, utilizing thresholds established at one-third and two-thirds as depicted in **Figure 4a**. Compared to the ES4^Hi^ group, the ES3^Hi^ group exhibited a significantly higher proportion of ES3 (*P* = 3 × 10^−4^, Kruskal-Wallis test) and a lower proportion of ES4 (*P* = 2 × 10^−2^, Kruskal-Wallis test) as illustrated in **Figure 4c**. We then examined processes associated with immune escape including clonal/subclonal neoantigen numbers, mutator phenotypes, TCR/BCR diversity, and Th1/Th2/Th17 signatures in the ES3^Hi^ and ES4^Hi^ groups. We observed that the ES3^Hi^ group exhibited a marked immune response, with increased neoantigen load, lower HRD score, reduced copy number burden, higher mutation rate, increased expression of Th1 cells, and higher T-cell receptor diversity scores compared to the ES4^Hi^ group (*P <* 0.01, Wilcoxon test), as depicted in **Figure 4b**. These findings suggest that the ES4^Hi^ group aligns with characteristics indicative of later stages of tumour evolution, displaying higher HRD and copy number burden, and having already developed mechanisms associated with immune escape.

### Evolutionary dynamics subtypes reflect driver mutations acting during late-stage cancer evolution

Based on our previous observations **(Figure 4)**, we hypothesize that the switch from an ES3^Hi^ state towards an ES4^Hi^ state during late-stage evolution, is accompanied by increased copy number burden and immune escape associated with dynamic changes in the tumour immune microenvironment. Mutation frequency generally displays an upward trend during clonal expansion under positive selection, while it tends to decrease when other stronger competitive subclone expand or under negative selection^13, 47, 48^. We posit that ESs capture groups of mutations with distinct growth behaviours during the evolutionary process of cancer. However, the specific mutation content within these groups may vary among individual patients. Therefore, identifying mutations undergoing the evolutionary process from an ES3^Hi^ state towards an ES4^Hi^ state can provide insights into the selection pressures acting upon driver mutations during late cancer evolution across various cancer types.

To detect the evolutionary modes of driver mutations, we constructed CCF distributions for the ES3^Hi^ and ES4^Hi^ groups separately for 409 consensus driver mutations identified by a comprehensive PanCancer analysis^49^. We then conducted a Kolmogorov-Smirnov test across cancer types to identify mutations with a significant transition between these two CCF distributions. The direction of the transition was determined by comparing the means of two CCF distributions. We define a driver mutation that acts on late-stage evolution in a type of cancer with a rightward transition from CCF distribution in ES3^Hi^ subtype to CCF distribution in ES4^Hi^ subtype. Such a transition of a driver mutation indicates the presence of stronger positive selection acting on subclones that do not carry this mutation. This observation implies a reduced significance of this driver mutation in driving switching between adaptive and innate immune mechanisms during late-stage evolution. Mutations show similar CCF distributions within ES3^Hi^ and ES4^Hi^ subtypes are considered neutral or early drivers that already reach fixation in both subtypes.

We found the CCFs distribution of several drivers in ES3^Hi^ move towards a higher frequency in ES4^Hi^ in most cancer types, notably ATRX, ARID1A, KMT2D, PTEN, ARID1A, PIK3CA, SETD2, KMT2C, RB1, CTNNB1, FAT1, FBXW7, BRAF, NOTCH1, CDKN2A, ERBB2 and KDM6A. For example, the CCFs distribution of FBXW7 in ES3^Hi^ (median = 0.71, n = 264) was significantly different from CCFs distribution in ES4^Hi^ (median = 1, n = 154) in colorectal cancer (*P* = 6 ×10^−8^, Kolmogorov-Smirnov test). Interestingly, we also observed a transition pattern toward lower frequency in CCF distribution between ES3^Hi^ and ES4^Hi^ for drivers in a few cancer types, which may be in line with negative selection. For example, the CCFs distribution of NF1 in ES3^Hi^ (median = 0.67, n = 180) was significantly different from the one in ES4^Hi^ (median = 0.29, n = 45) in BRCA (*P* = 1.5 × 10^−20^, Kolmogorov-Smirnov test) **(Figure 5a)**.

**Figure 5.**
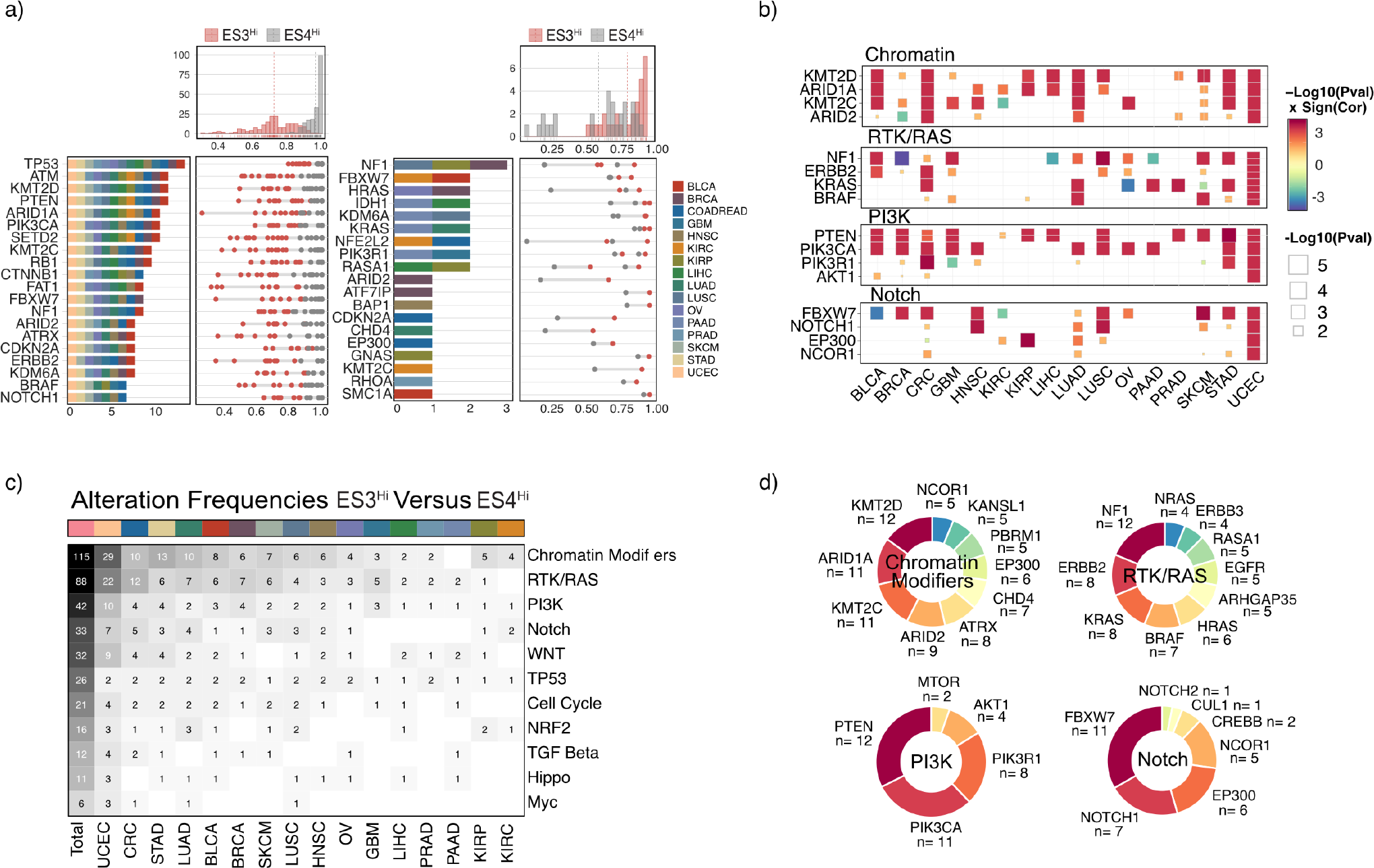
Evolutionary dynamics subtypes reflect different selection on driver mutations. **a**, Identification of mutations enriched in both ES3^Hi^ tumours and ES4^Hi^ tumours with a transition pattern on the CCF spectrum. **b**, Heatmap showing the distribution of mutations with identified transition in ES3^Hi^ versus ES4^Hi^ across cancer types and pathways. The colour spectrum shows the significance level and direction of identified transition on mutation CCFs distribution between ES3^Hi^ and ES4^Hi^ tumours. Only differences with false-discovery rate P < 0.05 (Kolmogorov-Smirnov test) are shown. **c**, Alteration frequencies distribution of enriched mutation with transition pattern across cancer types and pathways. **d**, Donut chart showing the frequency distribution of mutations identified with transition pattern per pathways.

Importantly, we found that clonal drivers under selection are enriched for key oncogenic pathways across cancer types, including chromatin, RTK/RAS, PI3K, and Notch **(Figure 5b)**. We found Chromatin Modifiers (including KMT2D, ARID1A, KMT2C etc.,), RTK/RAS (including NF1, ERBB2, KRAS, BRAF, etc.,) and PIK3 (including PTEN, PIK3CA, PIK3R1, etc.,) pathways account for a great part of the identified drivers under selection, especially in CRC, STAD, UCEC and LUAD **(Figure 5c,d)**. These observations suggest the roles of selection for specific drivers in different cancer types in regulating subclonal expansion and tumour-stroma interactions that drive switching between adaptive and innate immune mechanisms.

### Evolutionary dynamics subtypes show prognostic value in patient survival

We investigated further whether evolutionary dynamics subtypes are prognostic in 16 cancer types (Figure 6b). ES3^Hi^ and ES4^Hi^ samples were defined for each cancer type using one-quarter and three-quarters of the difference between ES4 and ES3 proportions as thresholds. We identified a worse progression-free survival for the ES4^Hi^ subtype in five cancer types, including CRC (HR = 2.5, 95%CI = (1.03,5.92), *P* = 0.043, log-rank test), KIRC (HR = 1.8, *P* = 0.05, log-rank test), LIHC (HR = 1.9, *P* = 0.043, log-rank test), PRAD (HR = 2.9, *P* = 0.004, log-rank test), UCEC (HR = 2.3, 95%CI = (1.11,4.98), *P* = 0.025, log-rank test) (Figure 6a).

**Figure 6.**
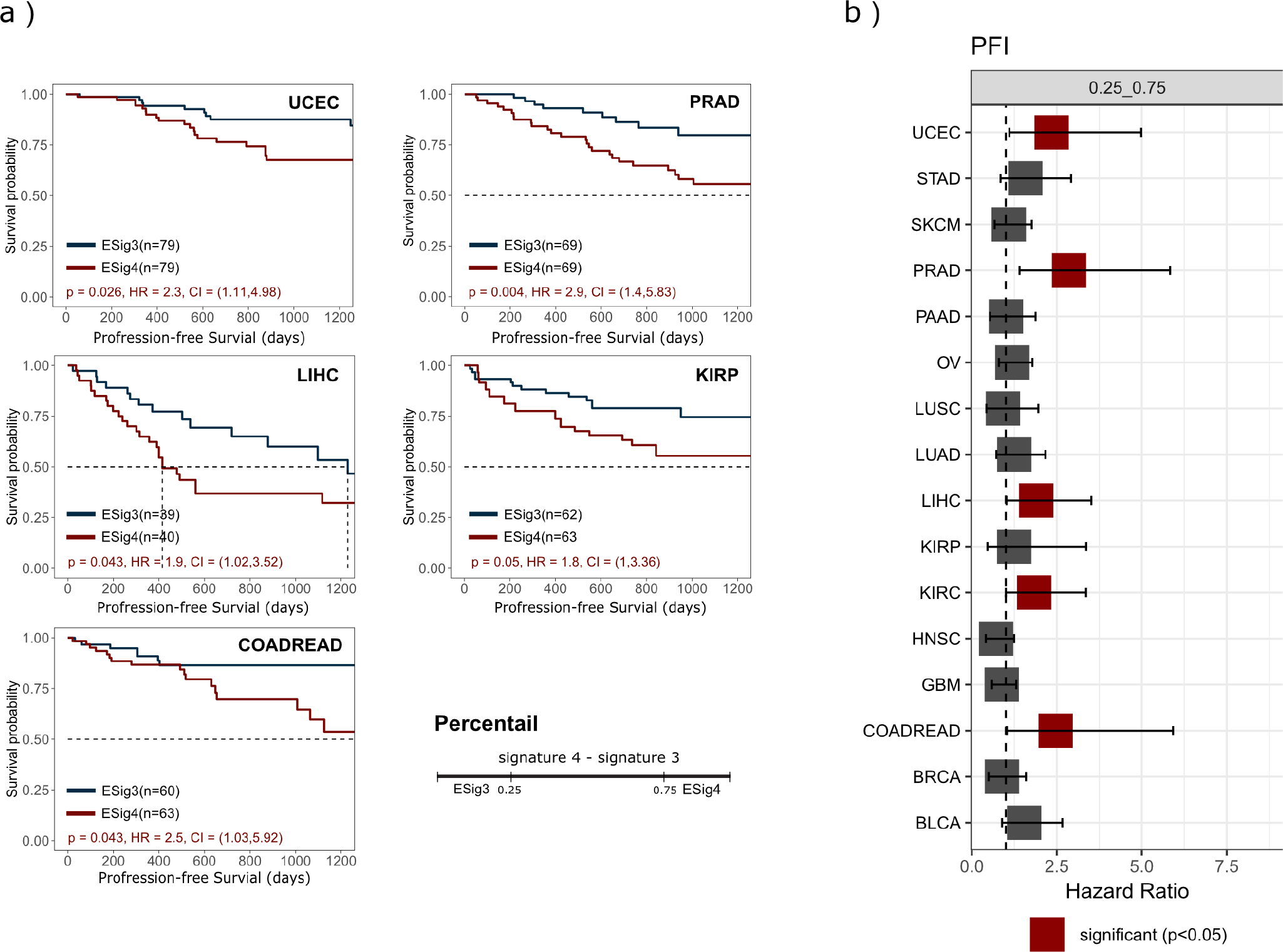
Evolutionary dynamics subtypes exhibit prognostic value. **a**, Kaplan-Meier plots with estimated hazards ratio and the 95% confidence interval show the difference in patient survival between evolutionary dynamics subtypes (using a quarter and three quarters as thresholds) in CRC, UCEC, LIHC, KIRP and PRAD. *P* values were computed from the Cox Proportional Hazards (CoxPH) regression modelling after applying a central log transformation to each signature exposure. The number of patients within each subtype is shown in the legend. **b**, Hazard ratio table for patient survival stratified based on ES subtype, derived from CoxPH regression model across cancer types.

## Discussion

In our pipeline, we identified consensus evolutionary signatures in tumours, termed evolutionary dynamics signature (ES), making assessments of evolutionary dynamics possible under the limitation of single time-point data for individual tumours. We approximate a generalised mathematical formula for modelling evolutionary dynamics in cancer genomes based on the population genetics theory as a data-driven solution using non-negative matrix factorization (NMF). Similar methods have been successfully applied to mutational signatures^26, 27^ and copy number signatures^28–30^. This framework consists of four major steps, (1) NMF-based type-wise signature extraction, (2) Hierarchical clustering into pan-cancer evolutionary dynamics signatures, (3) Signature assignment, and (4) Signature characterisation. These signatures can introduce interpretability by integrating with other patient-level data, such as gene expression, DNA damage scores, mutational signatures and immune infiltration. Besides, these signatures can be derived from bulk sequencing data like WES and WGS, which is rapid and cost-effective, providing great utility to clinical implementation.

Evolutionary signatures defined a continuum of cancer cell fractions representing neutral evolution, clonal expansion and fixation. Our analysis uncovered important pan-cancer correlations between evolutionary signatures and immune infiltration, DNA damage and cancer-driven mutational processes (Figure 2). Specifically, we identified a dynamic transition between adaptive and innate immune processes as tumours undergo clonal fixation and escape immune surveillance (Figure 3). tumours with high ES4 signature enrichment were associated with poor survival across several cancer types (Figure 6), highlighting the clinical utility of our approach. This work also reveals driver mutations that are specifically enriched during clonal expansion and fixation (Figure 5). The selection of distinct driver mutations in the context of lymphoid-poor and myeloid-rich immune micro-environments provides important insights into the dynamics of tumour progression across several cancer types.

We found that the quantification of ES contribution is influenced by sequencing coverage depth. Specifically, in the context of ES situated in regions of low CCF (ES1), the signature exposures estimated in both WGS and WES lack a robust correlation among individual patients (Figure 1e). A higher coverage level of sequencing data might be necessary to ensure the reliability of ES1 estimation. In this paper, we estimated the contribution of evolutionary dynamics signatures using a single bulk-sequencing sample for a patient. Further application to multi-region sequencing will be required to reflect the tumour spatial structure for individual patients. Our analysis does not provide prognosis implications using ES subtypes across all cancer types, suggesting tailoring cancer-specific signatures might be helpful to enhance the clinical application in specific cancer types.

In summary, evolutionary dynamics signatures provides valuable insight into how clonal expansion link to dynamic changes in the tumour immune microenvironment. We show that through signature analysis we can detect the clonal expansion of driver genes that modulate tumour-stroma interactions and identify subtypes with prognosis significance in many cancer types. Our study creates an opportunity to understand the complexity of ITH during cancer evolution and its potential implication in bulk-sequenced tumour samples.

## Data availability

PCAWG protected datasets are controlled access that is subject to data usage agreement. Somatic variant calls generated by PCAWG datasets is available for download at https://docs.icgc.org/pcawg/data/. In accordance with the data access policies of the TCGA projects, most molecular, clinical and specimen data are in an open tier which does not require access approval. Immune signatures used in this paper is described here^41^ and available for download. The source data underlying Figs. 2–6 and Supplementary Figs are provided as a Source Data file.

## Code availability

The code is available at Github repository.

## Acknowledgements

K.Y. acknowledges support from EP/R018634/1, BB/V016067/1, and European Union’s Horizon 2020 research and innovation programme under grant agreement No 101016851, project PANCAIM.

## Author contributions statement

X.Y, P.B., and K.Y. conceived the experiments. X.Y., W.L., and K.Y. conducted the experiments. G.M., P.V.L, F.M. conducted the copy number analysis. X.Y., P.B., and K.Y. analysed the results. X.Y., P.B., and K.Y. wrote the manuscript with feedback from all authors. P.B. and K.Y. jointly supervised the research.

## Additional information

To include, in this order: **Accession codes** (where applicable); **Competing interests** (mandatory statement).

The corresponding author is responsible for submitting a competing interests statement on behalf of all authors of the paper. This statement must be included in the submitted article file.

## Methods

### NMF-based signature extraction of cancer evolutionary dynamics in a tumour

We use non-negative matrix factorization (NMF) to solve this matrix factorisation problem and simultaneously extract the signatures and contributions of each signature among tumour samples. The framework of identification of evolutionary dynamics patterns based on NMF in this study performs separate steps as follows (see Figure1b):

### Step 1: Construct type-wise CCF-by-sample matrices

To assemble mutation size over CCF of tumours into matrices, the continuous interval of CCF between 0 and 1 is divided into 100 bins. The CCF-by-sample matrix was constructed for each cancer type. Each matrix has a size of 100 × *n*_*type*_, where *n*_*type*_ denotes the number of samples in a specific cancer type. In the CCF-by-sample matrix, columns represent tumour samples, rows represent the CCF span (100 rows as 100 intervals between 0 and 1), and each cell indicates the number of mutations falling within the corresponding CCF interval.

### Step 2: Determine the optimal number of signatures

A critical step of NMF is the estimate of factorization rank, i.e., the suggested number of signatures for factorization. Brunet algorithm^50^ was performed for 1000 runs with different random seeds between ranks 3-10. The rank was determined based on six quality measures (cophenetic coefficient, dispersion, evar, residual sum of squares, euclidean distance, and KL divergence). To detect overfitting, 1,000-time shuffles of the input matrix, by permuting the rows of each column, were also performed to get a null estimate of each of the scores. The rank was estimated for samples of different cancer types with the deemed optimal value under these constraints

### Step 3: Identification of consensus signatures of evolutionary dynamics

The normalised CCF-by-sample matrix of each cancer type was subjected to the NMF algorithm separately with the corresponding estimated rank in step 2. We normalise it here due to the huge difference in mutation burden among samples, ranging from tens to tens of thousands of mutations, which can make the evolutionary patterns of samples with small mutation loads obscure or even misclassified. Unsupervised hierarchical clustering was then performed on all signatures obtained to identify the consensus signatures across cancer types. The proper number of clusters was determined by the Hubert index as the significant peak in the Hubert index second difference plot. As a result, the consensus signatures obtained ended up averaging all the type-wise signatures allocated to the same cluster.

## Datasets

For each patient, we curated somatic mutation, integer-level copy number, tumour purity (fraction of tumour cells in the sample) and overall ploidy, donor clinical profiles and survival data from The Cancer Genome Atlas (TCGA). All somatic mutation samples from the TCGA were retrieved through the National Cancer Institute Genomics Data Commons Portal (TCGA Unified Ensemble "MC3"mutation calls, version 0.2.8). Only patients with matched germline (from blood samples) and primary tumour information available were considered.

### Estimation of cancer cell fractions

To estimate the cancer cell fraction (CCF) of somatic mutations in tumour samples, we used Ccube^33, 34^ algorithm, which allows for clustering and estimating cancer cell fractions (CCF) of somatic variants (SNVs/SVs) from bulk whole genome/exome data. The method takes the reference and alternative allele read counts of called variants, corrects for copy number alterations and purity, and then produces CCF estimates for all variants within the tumour sample. It identifies clusters of mutations, which can be used to determine the clonal architecture of individual tumour samples. Cancer cell fraction values larger than 1 (arising from sequence noise and copy neutral LOH events) were assumed to be 1.

### Signature assignment to individual patient samples

The consensus evolutionary dynamics signatures were used to assign an activity for each signature to 4146 TCGA patient samples. Linear Combination Decomposition (LCD) was performed to assign the amount of each signature harboured by tumour samples in terms of a decomposition of the given CCF-by-sample matrix *V* with known consensus signature *W* by solving the minimization problem *min*(∥*W* ∗*H* −*V* ∥) with additional constraints of non-negativity on H where W and V are known. After assigning signatures to each patient, we can estimate the contributions of each signature in the individual patient samples, which allows for subsequent patient-level analyses.

### Evolutionary dynamics signature validation

The signature identification procedure described above was applied to 2365 whole-genome sequenced samples from the ICGC Pan-Cancer Analysis of Whole Genomes Project (PCAWG). The number of signatures was fixed at 4 for matrix decomposition with NMF. Pearson correlation was computed between the TCGA signature-by-component weight matrix and the PCAWG signature-by-component matrix, signature by signature.

### Association with DNA damage, biological processes and immune microenvironment

We collected factors related to DNA damage from Thorsson et al.^41^, including homologous recombination deficiency (HRD)^42^, intratumor heterogeneity (ITH)^43^, aneuploidy Score^43^, copy number burden score ("Fraction Altered"and "Number of Segments")^42^, predicted neoantigen ("Indel Neoantigens"and "SNV neoantigens") and mutation rate^41^. Besides, we downloaded the weights of all known SBS mutational signatures for TCGA patients from COSMIC (v3.3 - June 2022).

To systematically investigate the interpretation of ESs in terms of the underlying immune microenvironment, we collected factors related to immune expression signatures, including macrophage regulation signature^51^ ("Macrophage Regulation"), immune cellular fraction estimates^41^ ("Macrophages"and "Monocytes"), lymphocyte infiltration^52^ ("Lymphocyte Infiltration Signature Score"), TGF-*β* response^53^ ("TGF-beta Response"), IFN-*γ* response^54^ ("IFN-gamma Response"), wound healing^55^ ("Wound Healing"), tumour-infiltrating lymphocytes from TCGA H&E images^41^ ("TIL Regional Fraction"), proliferation signature^54^ ("Proliferation"), leukocyte and stromal fractions^41^ ("Leukocyte Fraction"and "Stromal Fraction"). we also collected factors related to immune mechanisms, including immune escape annotation^56^, clonal and subclonal neoantigen numbers^56^, mutator phenotypes^57^, T-cell receptor (TCR) and B-cell receptor (BCR) diversity^41^, Th1/Th2/Th17 signatures^57^.

We evaluated the association of these factors and the constituent components of evolutionary dynamics signatures for 4146 TCGA patients across cancer types. Association between evolutionary dynamics signature exposures and features related to DNA damage, biological processes and immune microenvironment were performed using one of two procedures: for a continuous association feature, Spearman correlation was performed with adjusted p values for multiple testing using the Benjamini-Hochberg method^58^; for a binary association feature, samples were divided two groups and a Mann-Whitney U-test was performed to test for differences in signature exposure medians between groups. Besides, the association between evolutionary dynamics subtypes and features related to DNA damage, biological processes and microenvironment were performed using the Wilcoxon signed-rank test. The Kruskal-Wallis test was also performed to test the differences between groups.

### Identification of evolutionary dynamics subtypes

To investigate the various states of evolutionary dynamics characterized by distinct immune mechanisms, we categorised tumours into ESig3^Hi^ or ESig4^Hi^ subtypes, separately for each cancer type. This subtyping was based on the disparities in proportions of ES3 and ES4, using predefined thresholds set at one-third and two-thirds. Consequently, the ESig3^Hi^ and ESig4^Hi^ subtypes were established as balanced labels for each specific cancer type, enabling the exploration of group-wise differences in immune mechanisms and prognosis.

### Detection of selection modes on driver mutations in cancer

We constructed CCF distributions in ES3^Hi^ and ES4^Hi^ groups separately for 409 cancer consensus driver mutations identified by a PanCancer analysis^49^ to detect the evolutionary modes of driver mutations. We then identified the mutations with a significant transition between these two CCFs distributions using the Kolmogorov-Smirnov test across cancer types. The transition direction was determined based on the mean of two CCFs distributions. We define a driver mutation that undergoes positive selection in a type of cancer with a rightward transition from CCF distribution in ES3^Hi^ subtype to CCF distribution in ES4^Hi^ subtype. Driver mutations with opposite transition directions are determined as under negative selection. Mutations show similar CCF distributions within ES3^Hi^ and ES4^Hi^ subtypes are considered as neutral or early drivers that already reach fixation in both subtypes.

### Survival Analysis

Cox Proportional Hazards (CoxPH) regression modelling was used to determine whether ES subtype (ES3^Hi^ and ES4^Hi^) predicts patient survival. A central log transformation was applied to each signature’s exposure prior to its submission to the CoxPH model. The Hazard Ratio (HR) and the 95% confidence interval (95%CI) of HR were calculated with p values. A False Discovery Rate (FDR) correction using the BH method was applied to p values. A test of Schoenfeld residuals was performed to assess the PH assumption. The Kaplan-Mier estimator was used to create the survival plots and the log-rank test was used to compare the difference in survival curves.

## Supplementary Materials

### Rank estimate for NMF on TCGA cohorts

To extract type-wise evolutionary dynamics signatures using NMF for the TCGA cohort, we need to determine the proper rank for each cancer type separately following the above rank estimate procedure.

To achieve this, we run 1000 runs of NMF with Brunet’s algorithm for each rank between 3 and 12 and each cancer type using the original and randomised datasets. We then compute seven quality measures for each condition, including sparseness, residual sum of squares, explained variance, dispersion coefficient, cophenetic correlation coefficient, euclidean distance and Kullbakc-Leibler divergence (Supplementary Figure 1).

We choose the best value of factorisation rank for each cancer type suggested by the three mentioned methods and finally combine the three suggestions and inspection checks to determine the final chosen rank (**Table 1**). As a result, we performed 1000 runs of NMF with Brunet’s algorithm based on the chosen rank for each cancer type and obtained the type-wise evolutionary dynamics signatures (**Supplementary Figure 2**).

**Supplementary Figure 1.**
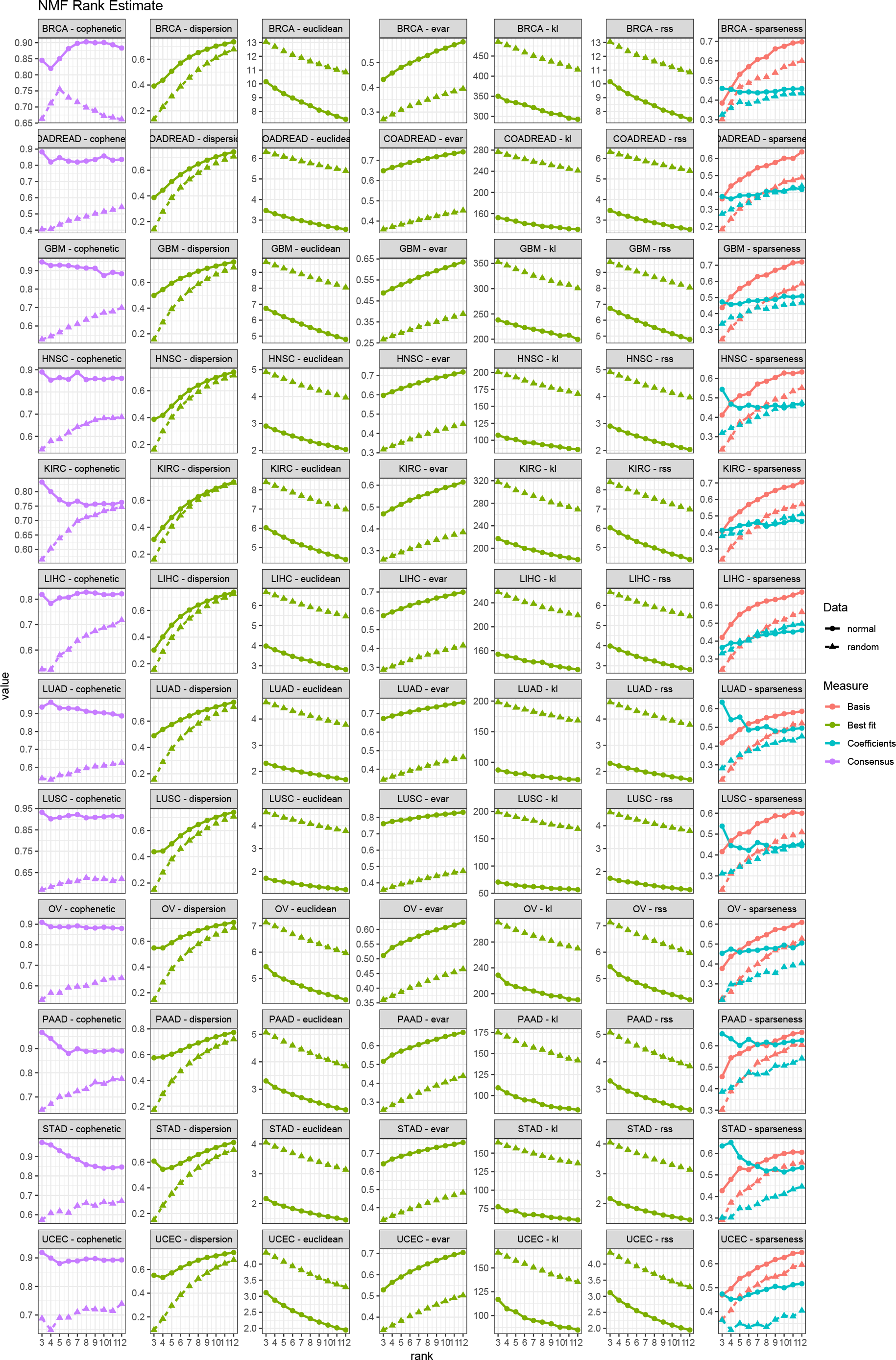
Estimation of the NMF rank across 12 types in TCGA cohort: comparison of quality measures computed for each rank value between 3-12. Each point on the graph was obtained from 1000 runs of NMF with Brunet’s algorithm. The curves for the actual data are in a circle shape, and those for the randomized data

**Supplementary Table 1.** Suggested rank for TCGA cohorts.

| Suggested rank for TCGA cohorts |  |  |  |  |
| --- | --- | --- | --- | --- |
| Type | brunet et al. | hutchins et al. | frigyesi et al. | Chosen rank |
| BRCA | 5 | 6 | 10 | 6 |
| COADREAD | 3 | 5 | 7 | 5 |
| GBM | 9 | 6 | 12 | 6 |
| HNSC | 3 | 5 | 5 | 4 |
| KIRC | 3 | 5 | 9 | 5 |
| LIHC | 4 | 4 | 6 | 4 |
| LUAD | 4 | 5 | 4 | 4 |
| LUSC | 3 | 4 | 4 | 4 |
| OV | 3 | 4 | 6 | 5 |
| PAAD | 4 | 4 | 5 | 5 |
| STAD | 4 | 4 | 5 | 4 |
| UCEC | 4 | 4 | 9 | 4 |

### Rank estimate for NMF on PCAWG cohorts

To extract type-wise evolutionary dynamics signatures using NMF for the PCAWG cohort as validation, we performed the rank estimate procedure similar to TCGA by running 1000 runs of NMF with Brunet’s algorithm and computing quality measures for each rank between 3 and 12 and each cancer type using the original and randomised datasets (**Supplementary Figure 3**). We choose the best value of factorisation rank for each cancer type suggested by the three mentioned methods and finally combine the three suggestions and inspection checks to determine the final chosen rank (**Table 2**). As a result, we performed 1000 runs of NMF with Brunet’s algorithm based on the chosen rank for each cancer type and obtained the type-wise evolutionary dynamics signatures (**Supplementary Figure 2**).

**Supplementary Table 2.** Suggested rank for ICGC cohorts.

| Suggested rank for ICGC cohorts |  |  |  |  |
| --- | --- | --- | --- | --- |
| Type | brunet et al. | hutchins et al. | frigyesi et al. | Chosen rank |
| BRCA | 3 | 5 | 4 | 3 |
| CLLE | 3 | 4 | 4 | 3 |
| ESAD | 4 | 4 | 4 | 4 |
| LIRI | 4 | 5 | 4 | 4 |
| MALY | 5 | 5 | 4 | 5 |
| OV | 3 | 4 | 4 | 4 |
| PACA | 4 | 4 | 4 | 4 |
| PBCA | 3 | 4 | 4 | 4 |
| PRAD | 3 | 4 | 4 | 4 |

**Supplementary Figure 2.**
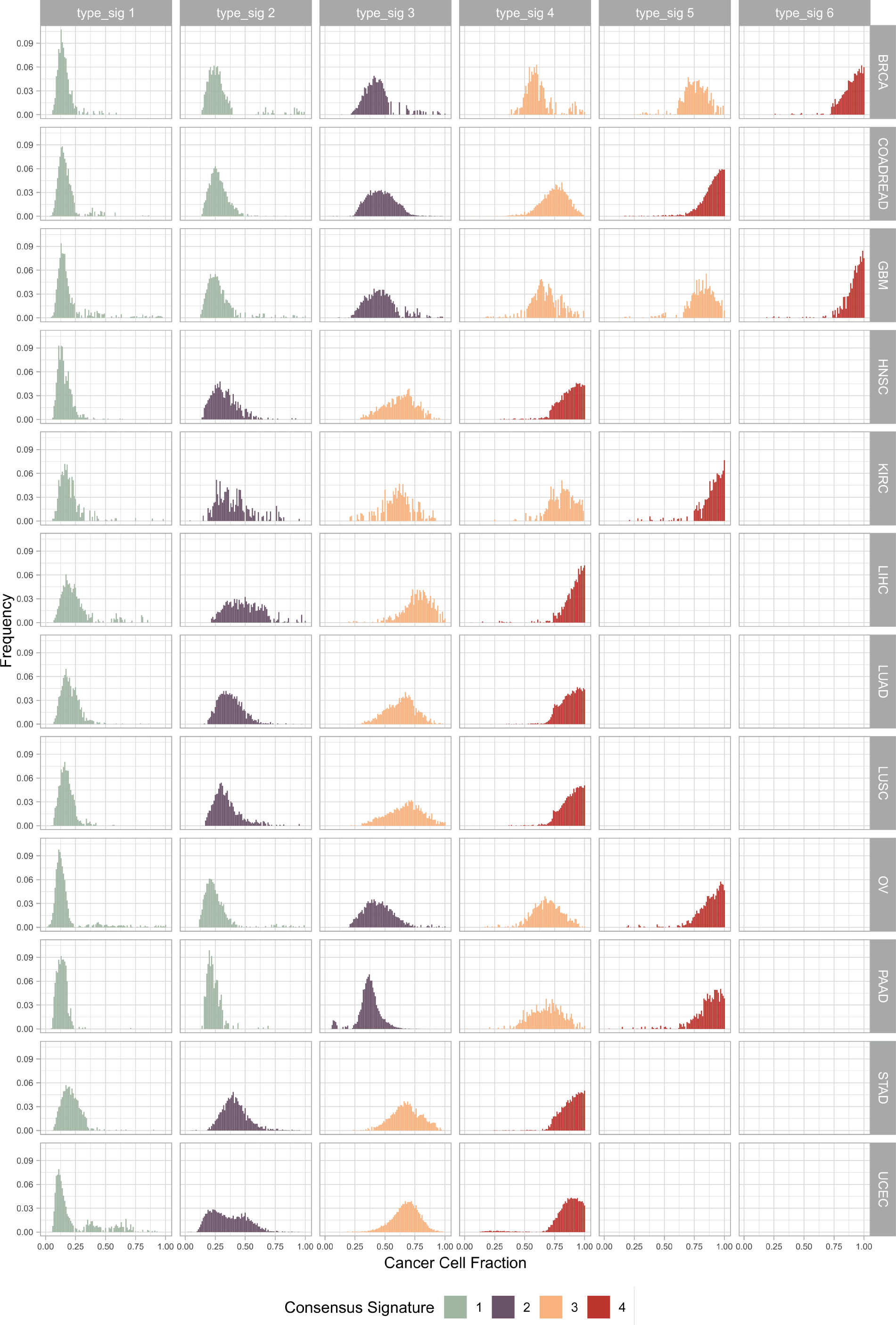
The evolutionary dynamics signatures obtained for each cancer type in TCGA following the suggested ranks (Supplementary Table 1). Colour represents the assignment results with the following hierarchical clustering for ESs shown in Figure 1.

**Supplementary Figure 3.**
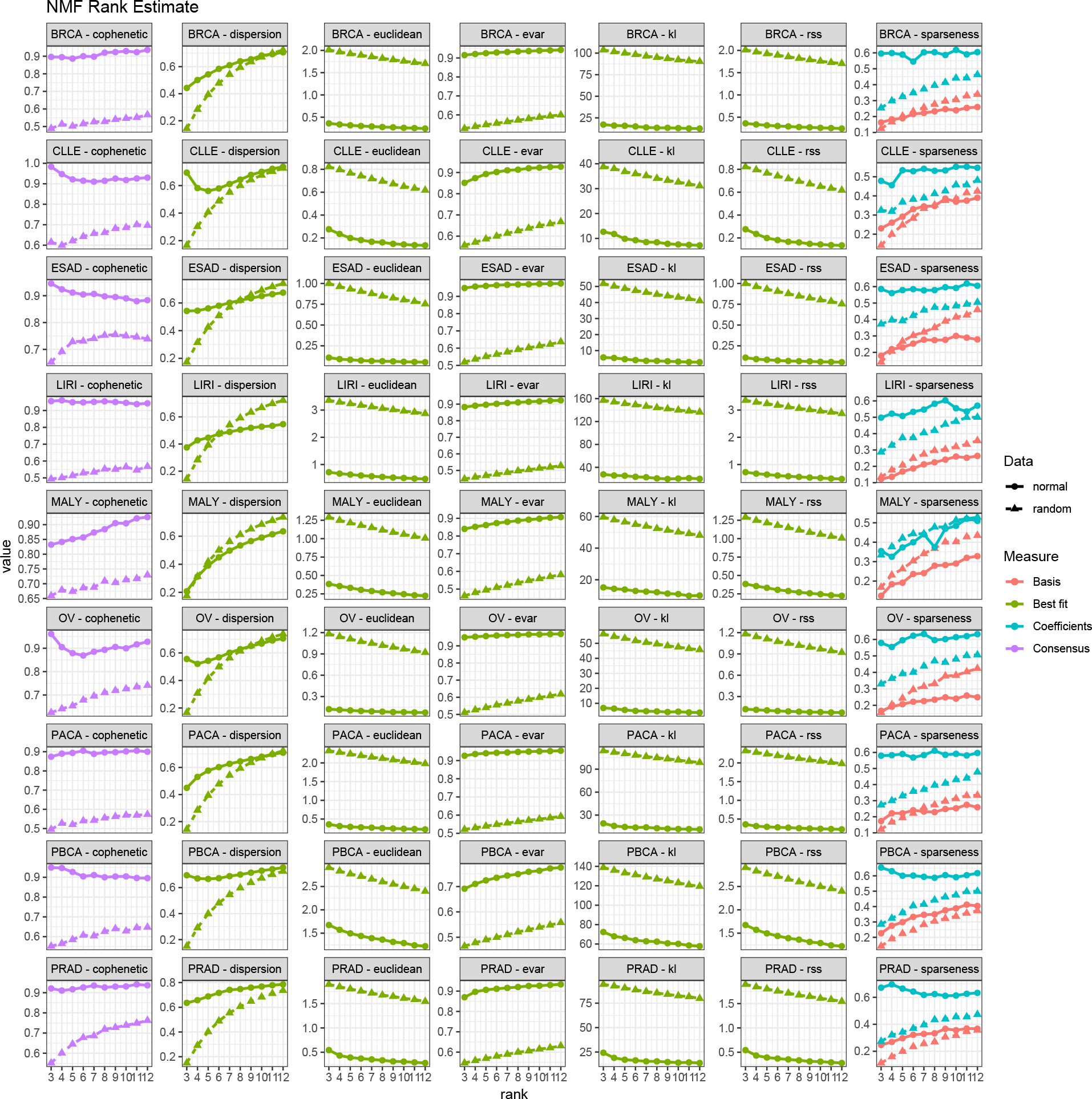
Estimation of the NMF rank across 9 types in PCAWG cohort: comparison of quality measures computed for each rank value between 3-12. Each point on the graph was obtained from 1000 runs of NMF with Brunet’s algorithm. The curves for the actual data are in a circle shape, and those for the randomized data are in a triangle shape.

